# Transfer learning and DNA language models enhance transcription factor binding predictions

**DOI:** 10.1101/2024.11.08.622635

**Authors:** Ekin Deniz Aksu, Martin Vingron

**Affiliations:** Department of Computational Molecular Biology, Max Planck Institute for Molecular Genetics, Ihnestraße 63-73, 14195 Berlin, Germany

## Abstract

Identification of *in vivo* transcription factor (TF) binding sites is crucial to understand gene regulatory networks, but the lack of scalability in the methods for their experimental identification directs researchers towards computational models. TF binding site prediction models are often specific for a given TF, which also hinders the generalizability of models to previously unseen TFs. Here, we present an approach to predict *in vivo* TF binding sites using DNA accessibility, TF RNA expression and TF binding motifs. Our novel method leverages DNA language model embeddings and transfer learning to improve its accuracy and generalizability, achieving a mean area under the precision-recall curve (AUPR) of 0.51 in held-out cell types and chromosomes in the ENCODE-DREAM *in vivo* TFBS prediction challenge, outperforming the top-ranked methods. Furthermore, we show that prediction accuracy increases when TFs are highly active and exhibit cell-type specific expression. We finally test our models in an independent dataset on previously unseen TFs, and report a mean AUPR of 0.36, which is state-of-the-art in a cross-TF, cross-cell type and cross-chromosomal setting.

## Introduction

Gene expression is regulated by a set of transcription factors (TFs) that bind to regulatory regions on the DNA^1^. Although sequence-specific binding preferences are known for many TFs, only a subset of predicted TF binding sites (TFBS) are occupied in any given cellular context —referred to as *in vivo* TFBS. Accurate identification of *in vivo* TFBS requires chromatin immunoprecipitation assays such as ChIP-seq^2^, which is impractical to perform across all 1600 human TFs due to their dependency on specific antibodies and the lack of parallelization or multiplexing.

Computational methods have emerged as powerful alternatives for predicting *in vivo* TFBS without the need for exhaustive experimentation. These methods broadly fall into three categories: i) supervised tools that use motif position weight matrices (PWMs) along with DNase-seq, ATAC-seq and/or histone modification data, such as Virtual ChIP-seq, FactorNet, maxATAC and others^3–12^. These tools are trained on TF ChIP-seq data and can make whole-genome TF binding predictions on unseen tissues from the provided data. ii) Tools that are based on PWM-scanning in accessible regions, such as TF footprinting (HINT-ATAC^13^, TOBIAS^14^) or gene regulatory network (GRN) identification methods such as SCENIC+, Inferelator, GRaNIE, and most recently LINGER^15–19^. These methods are not trained on ChIP-seq data, and they usually predict TF binding in gene promoters, enhancers or ATAC-seq peaks as part of GRN analysis. iii) Sequence-to-activity deep learning models such as DeepBind, Enformer and others^20–23^. These methods learn the “complex grammar” of TF cooperation, but since they take only DNA sequence as input, they are limited to predictions on cell types in the training set.

Training supervised models requires high-quality ChIP-seq labels for each TF, which are not readily available for all TFs. Moreover, prediction accuracy is improved when data from multiple cell types are incorporated^12^. Therefore, efficient integration of multiple data sources and building prediction models that are generalizable to unseen TFs remain key objectives. We propose leveraging two emerging technologies to achieve this: transfer learning and DNA language models. Transfer learning enhances prediction accuracy by using models that are pre-trained on other datasets, thus offering a way to integrate data efficiently. On the other hand, DNA language models such as DNABERT^24^ and the Nucleotide Transformer^25^ extract high-level abstract features from DNA sequences, and can potentially improve generalizability because they learn universal rules regarding genomes.

In this study, we present an approach to predict *in vivo* TFBS using DNA accessibility, TF RNA expression and TF binding motif information. We extracted a set of handcrafted, biologically informed features from DNA regions and supplemented them with embeddings from DNA language models. Using these features, we trained gradient-boosted decision tree models on ground-truth TF ChIP-seq data from the ENCODE-DREAM challenge and compared prediction performance to the top-ranked challenge contestants.

We employed a novel two-stage transfer learning method that leverages all available data efficiently. First we built a general prediction model which is trained on all training TFs and cell types. Secondly, we trained new TF-specific models for each individual TF, while incorporating the general model’s predictions as an additional input feature. We prioritized model interpretability and investigated the contributions of various biological features to prediction accuracy. Additionally, we explored the factors driving variation in prediction performance across different TFs, and show that TFs with tissue-specific expression profiles are predicted more accurately than constitutively expressed TFs.

Finally, we tested the performance of our models in previously unseen TFs, in a very challenging cross-TF, cross-tissue, and cross-chromosome setting. We show that our general model outperforms the mean TF-specific model ensemble in cross-TF predictions. Remarkably, this model maintains a strong predictive power even under these challenging conditions.

## Results

### Model overview and training scheme

The task is to predict TF binding probability in regulatory regions given a TF and a cell type. To achieve this, our model incorporates handcrafted features that fall into four categories: i) genomic features which depend only on the genomic region, ii) motif features which depend on the region and the TF in question, iii) cell type features which vary based on the region and the cell type, and iv) cell-and-TF features which depend on both the TF and the cell type (Fig. 1a). A detailed description of all features and their definitions is provided in the Methods section.

**Fig 1:**
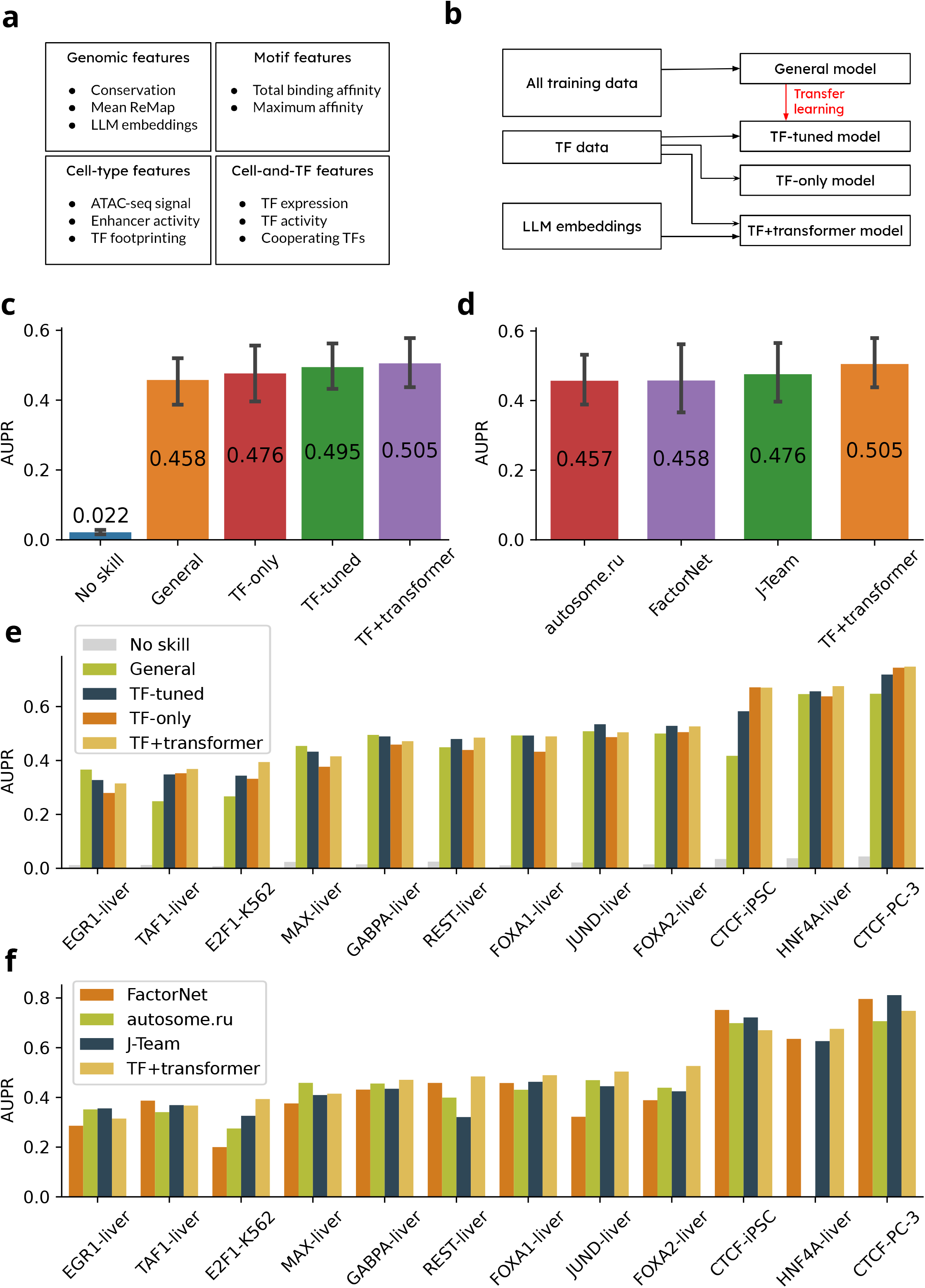
Accurate prediction of cell type specific in vivo TFBS. a)Overview of features used in the models. b) Schematic of the model training process. The transfer learning approach is implemented by incorporating the general model’s prediction as an additional feature when training the TF-tuned model. c, d) Area under the precision-recall curve (AUPR) comparing the performance of our models and against challenge winners (n=12). e, f) AUPR scores across TFs.

We developed four different types of gradient-boosted decision tree models. First, a “general model” was built, which pools all available training data across TFs and cell types, allowing the model to generalize across different conditions. Each genomic region is included multiple times in the training set, as many as the number of different TF-cell type pairs. This significantly expands the training set which is beneficial for learning optimal parameters. Second, we used transfer learning to develop “TF-tuned” models, which are fine-tuned to predict binding of individual TFs. The fine-tuning is achieved by incorporating the general model predictions as an additional input feature. Third, we developed “TF-only” models which only use training data from the selected TF, using the same feature set as the general model. Lastly, our “TF+transformer models” use, in addition to the handcrafted features, 1024-dimensional embeddings from the DNA language model Nucleotide Transformer^25^ (Fig 1b).

Our training data comes from three main sources: ATAC-seq and RNA-seq data from the ENCODE project^26^ and TF binding motifs from the TRANSFAC^27^ database. We further utilize precomputed^28^ enhancer probabilities derived from histone modification ChIP-seq data using the CRUP tool^29^. Training genome regions were defined as all ENCODE candidate cis-regulatory elements (cCREs)^26^, excluding regions on chromosomes 1,8, and 20, which were reserved for testing. Altogether, our dataset contains 8 cell types, 29 TFs and 75 TF-tissue pairings.

### Accurate prediction of TF binding sites

We evaluated the performance of our models on 12 TF-cell type pairs, corresponding to the “final round” dataset of the ENCODE-DREAM challenge. For each test region, we compared the model prediction of TF binding to the ground truth TF ChIP-seq labels provided by the challenge. Our evaluation metric is the area under the precision-recall curve (AUPR), chosen due to the significant class imbalance in the dataset, where bound labels represent only ∼1% of samples. In contrast, the area under the receiver-operating characteristic (ROC) curve fails to differentiate between strong and weak models (Supplementary Fig. 1).

All four of our models show very good overall performance with mean AUPR scores ranging between 0.46 and 0.505. The mean AUPR for random guessing is 0.02, which corresponds to the ratio of TF-bound enhancers in the test set. The TF+transformer model surpassed the other models with an AUPR of 0.505, compared to TF-tuned (0.495), TF-only (0.476) and the general model (0.458) (Fig 1c). Interestingly, even though the general model is fixed and does not change depending on the tested TF, it had a comparable performance to TF-specific models.

Remarkably, even though the general and TF-tuned models used the same training data, transfer learning lets the TF-tuned models adapt to the intricacies of different TFs and thus improve their performance. TF-tuned models are also more accurate compared to traditional TF-only models, underlining the benefits gained from integration of data from multiple TFs.

Performances of these models changed substantially based on the chosen TF, with AUPR scores ranging from 0.31 to 0.75. The highest score was obtained for CTCF, which is easier to predict than most TFs probably due to its high motif information content and constitutive binding across different cell types. Lowest scores are obtained for EGR1 and TAF1 in liver cells, followed by E2F1 in the K562 cell line (Fig 1e). Interestingly, there were some TFs where performance deviated from the general trend. For example, in EGR1, GABPA and MAX, the general model had the best performance. Tuning the general model further in these cases proved detrimental to performance. The reason behind this may be lower data quality in these TFs compared to other datasets.

Close examination of a single decision tree from the TF-tuned FOXA2 model reveals transfer learning in effect. The tree initially checks the value of the general model score. If this score is close to zero, no additional features are queried and the prediction is steered towards an unbound state. Conversely, a very high prediction score pushes the prediction towards a bound state. These represent cases where the general model has high confidence. In contrast, if the general model has low confidence, represented by an intermediate score, the tree proceeds to evaluate additional features such as enhancer activity or co-operating TFBS, thus further refining its prediction (Fig. 2).

**Fig 2:**
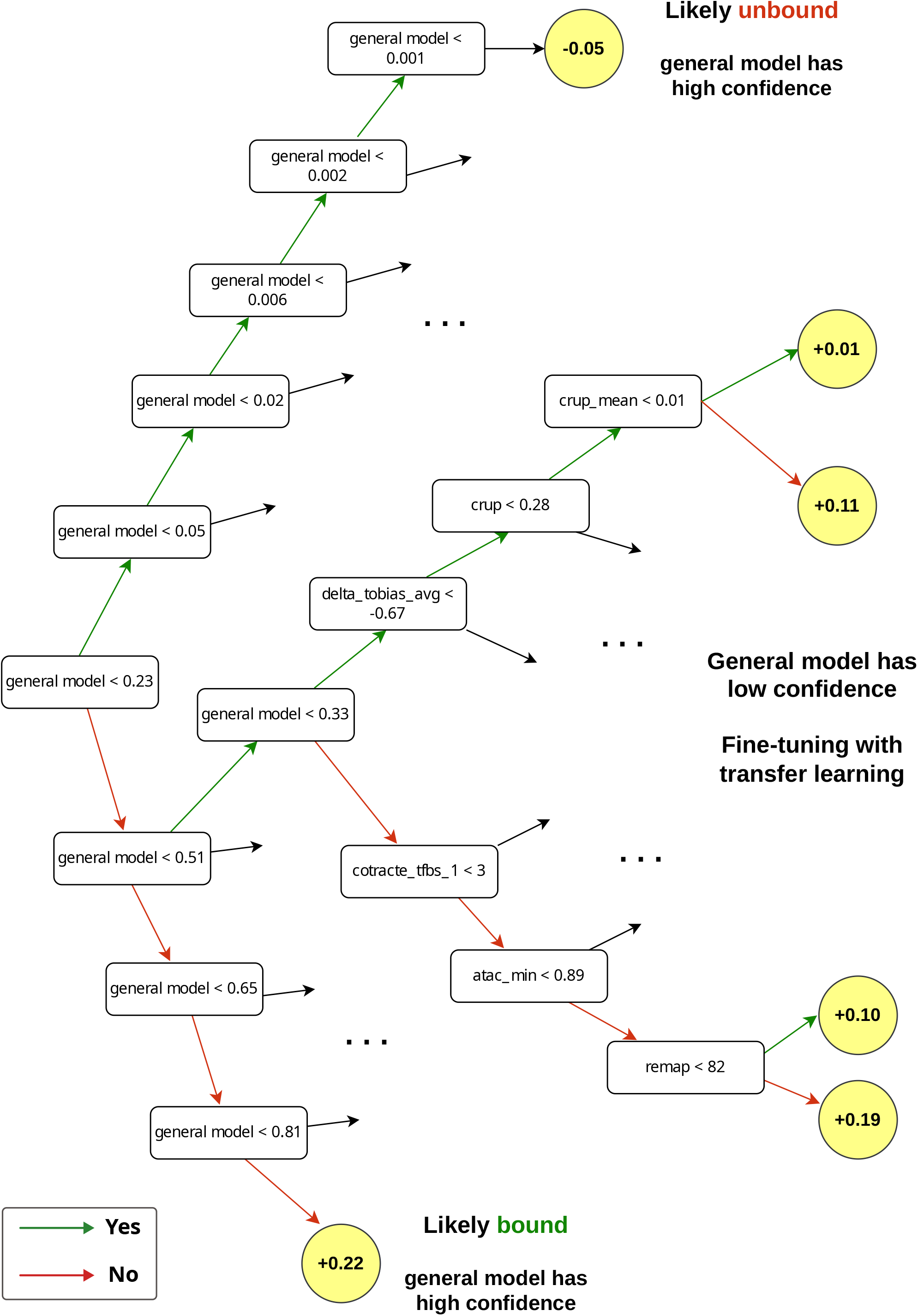
Transfer learning-informed decision trees. An excerpt from a representative decision tree from the FOXA2 TF-tuned model. Nodes with a white background represent decision splits based on feature values, while yellow-background nodes correspond to output leaves indicating predictions. Top and bottom paths denote paths where the general model has high confidence, and hence the model does not probe other features. In the middle of the tree, the general model score is intermediate, and fine-tuning is achieved by probing other features in a TF-specific manner.

### Comparison to other methods

We have selected three top-performing methods from the ENCODE-DREAM challenge for comparison: autosome.ru (https://www.synapse.org/Synapse:syn8006653/), FactorNet^9^ and J-Team^11^, with the latter tied for first place.

Our TF+transformer model’s mean AUPR of 0.505 was the highest, followed by J-Team (0.48), FactorNet (0.46) and autosome.ru (0.46) (Fig 1d). Remarkably, even our general model performed comparably to autosome.ru and FactorNet, while the TF-tuned and TF+transformer models were further ahead.

Since the other methods did not have access to histone modification and ReMap data, we also tested a “Dream-like” setup to assess the impact of additional data sources. We omitted these data types and retrained new models, and our models were still ahead despite this stricter condition, showing only a modest decrease of 0.02 AUPR on average. It is important to highlight that these tools had a much larger context window spanning multiple kilobases, which is advantageous for performance. In contrast, our models were deliberately designed with a constrained context window of one enhancer (max. 350 bp), prioritizing interpretability over performance. A more detailed examination of method comparability is provided in the Supplementary Information.

Examining the performances in specific TFs, our model performed consistently and was ranked as the best predictor in 7 out of the 12 selected TF-tissue pairs. However, CTCF stood out as an exception, where FactorNet and J-Team had a lead in performance. This is most likely because the information-rich CTCF motif makes sequence-based features more critical for accurate prediction (Fig. 3e). Given that FactorNet is a convolutional neural network with direct sequence input, it is better suited to capture higher level sequence features.

**Fig 3:**
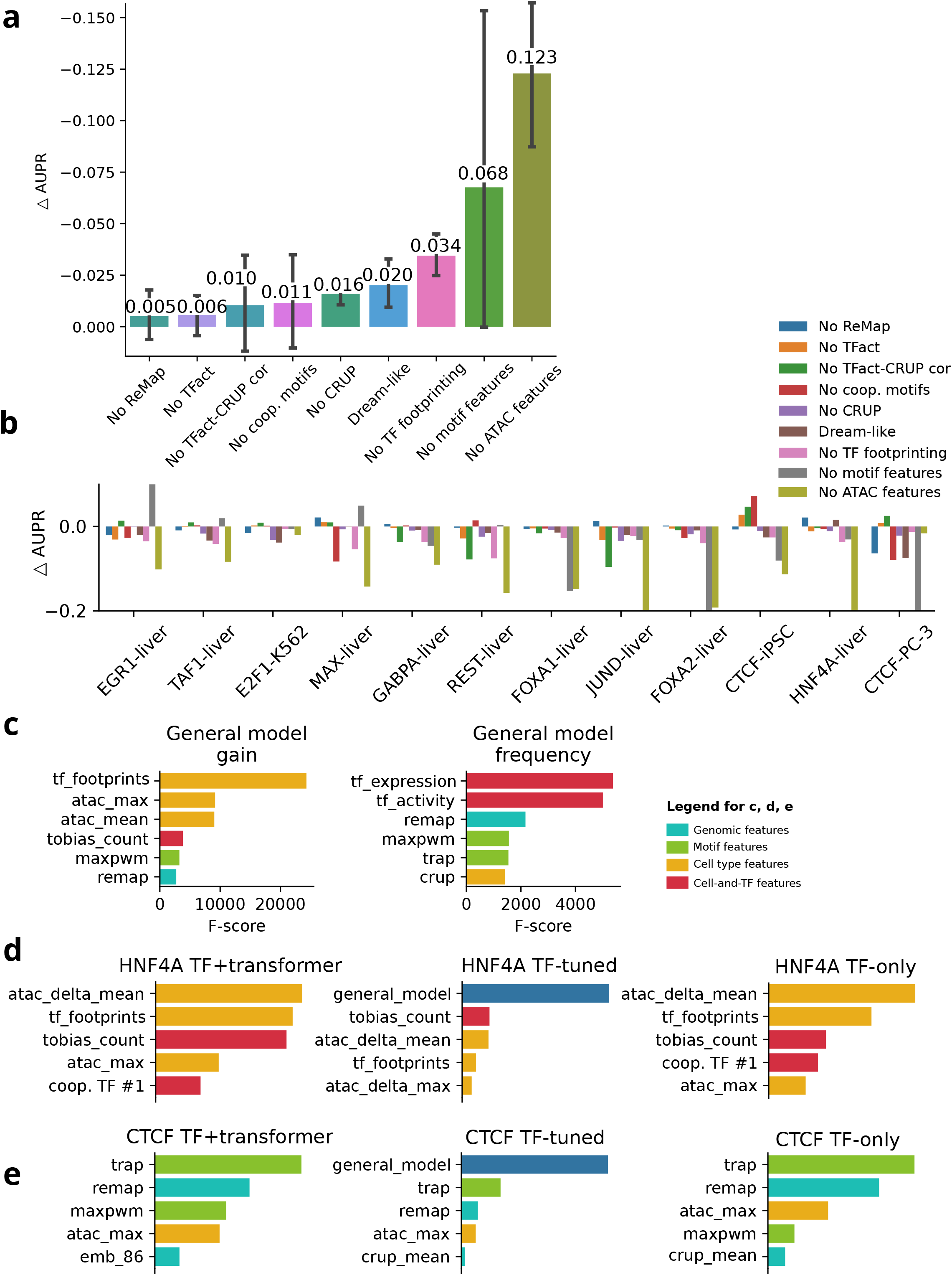
Feature importance. a)The change in AUPR (ΔAUPR) compared to the full TF-tuned model after masking specific feature sets. Error bars indicate variance across TFs, as shown in (b) (n=12). c) Feature importance ranking based on XGBoost F1-scores, with gain and frequency metrics. d, e) Feature contributions in different HNF4A and CTCF models. Features are colored according to their class.

### Impact of different features

We systematically tested the contribution of different features to model performance, by omitting specific feature sets from the training set and retraining general and TF-tuned models. Since many of our features are interdependent, we omitted subsets of features as opposed to single features. For example, motif-based features were excluded together as they are highly correlated.

On average, ATAC-seq-derived features were the most impactful, causing a notable AUPR drop of 0.12, which represented 24% of the base performance. Motif-based features followed with a decrease of 0.07 AUPR. In contrast, removing other features such as CRUP enhancer scores, TF activity or ReMap scores had minimal impact, with a ΔAUPR of 0.02 or below.

ATAC-seq features were the most important in 8 of the 12 TFs tested, consistently making significant contributions, with the exception of E2F1 in K562 cells and CTCF in PC-3 cells. On the other hand, motif-based features showed a more complex contribution pattern. For TFs such as EGR1, TAF1, HNF4A, and MAX, omitting these features slightly improved performance. TF activity-CRUP score correlations were important in the GABPA, REST and JUND models, but they are detrimental to performance in CTCF (Fig 3b).

A closer look at individual features revealed that the TF footprinting score stands out as the single most influential feature in the general model (Fig. 3c). TF footprinting incorporates both accessibility and binding affinity, while accounting for local decreases in accessibility due to TF binding. Therefore, it captures a high degree of information regarding TF binding events. Also, when analyzing feature importance using XGBoost, we found that it is important to choose the correct metric: the “frequency” metric is misleading because it simply counts how often a feature is included in decision tree splits, and less important features are sometimes included more often.

The importance of specific features also varied by TF and model architecture. For HNF4A, features such as ΔATAC, TF footprinting and bound TFBS counts are important in TF+transformer and TF-only models. In the TF-tuned model, expectedly, the general model score was by far the most important feature (Fig. 3d). Interestingly, for the CTCF TF+transformer model, sequence-based features were in the lead, with total motif affinity being the most important. One of the DNA language model embedding dimensions was also in the list of top features. It is possible that this dimension captured some sequence feature regarding CTCF binding (Fig. 3e).

### Prediction performance on HNF4A in example locus

We analyzed the HNF4A ChIP-seq signal in liver tissue in different genomic regions, comparing 3000 randomly selected enhancers to the top 3000 enhancers ranked by various features (Fig. 4a). Regions ranked by the highest correlation between TF activity and CRUP scores show a signal in nearly half of the regions, centered around selected enhancers. The signal intensity increased when using CRUP scores or ATAC-seq signals. Ranking enhancers by TFBS score results in a clear and strong signal from the top-ranked enhancers but the signal drops off faster compared to ATAC-ordered regions. Using the number of TF footprinting-predicted bound TFBS gives a better result. The best result is achieved by the TF+transformer model, where a strong signal centered around the enhancer is present in almost all of the top 3000 predictions (Fig 4a).

**Fig 4:**
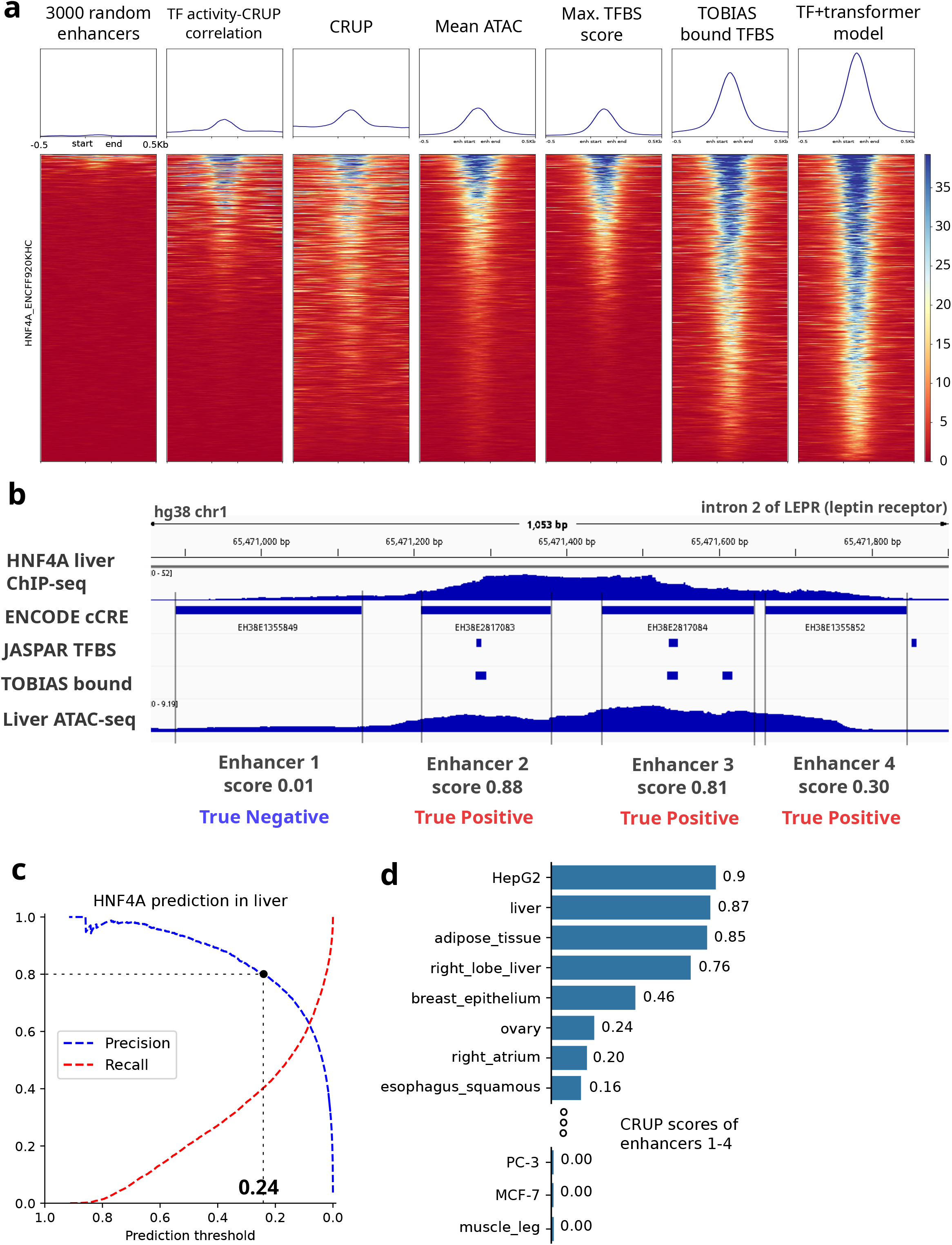
Epigenomic signals from selected enhancers. a)HNF4A ChIP-seq signal (fold change compared to control) from liver cells, centered on enhancers ranked by various metrics. b) ChIP-seq, ATAC-seq signals, and TFBS from four example enhancers located in intron 2 of the LEPR gene. c) Precision and recall curves for HNF4A in liver tissue, d) Mean CRUP enhancer scores for the selected enhancers.

To assess prediction accuracy at a specific locus in the human genome, we examined four enhancers located in intron 2 of the *LEPR* (leptin receptor) gene, which is linked to feeding response and likely to be active in liver tissue. This locus is accessible in liver, with HNF4A binding sites present in enhancers 2 and 3. Our TF+transformer model predicts HNF4A binding probabilities of 0.01, 0.88, 0.81, and 0.3 for enhancers 1 through 4, respectively. Overlaying with the HNF4A ChIP-seq signal, the predictions align well with the observed binding pattern (Fig. 4b). Precision and recall analyses indicated that the threshold for 80% precision is around 0.24, confirming enhancers number 2, 3 and 4 as true positives, while enhancer 1 is a true negative (Fig. 4c).

Enhancer activity profiles of the selected enhancers show that top ranked biological states are relevant to liver, including HepG2 (hepatocellular carcinoma cell line), liver, adipose tissue and right lobe of liver, all showing high enhancer probabilities. In contrast, enhancer activity drops off sharply in other tissues, many with CRUP scores of zero. This pattern is consistent with the observation that these enhancers are likely liver-specific.

### Prediction performance indicators

To explore potential predictors of model performance, we examined correlations between performance and TF-related indicators such as mRNA expression levels and TF activity. TF activity is defined as the contribution of each motif to cell type-specific enhancer activity (see Methods). For each TF-tissue pair, we plotted model performance against TF expression in the relevant tissue. We included TFs from the leaderboard set of the ENCODE-DREAM challenge to boost sample size, and we excluded CTCF because its exceptionally constitutive binding makes it much easier to predict than other TFs. We observed no statistically significant linear relationship between model performance and TF expression. However, when TF activity was substituted for expression, a statistically significant relationship emerged (Pearson’s correlation, p < 0.05), indicating that TF activity is a better predictor of performance (Figures 5a, b).

**Fig 5:**
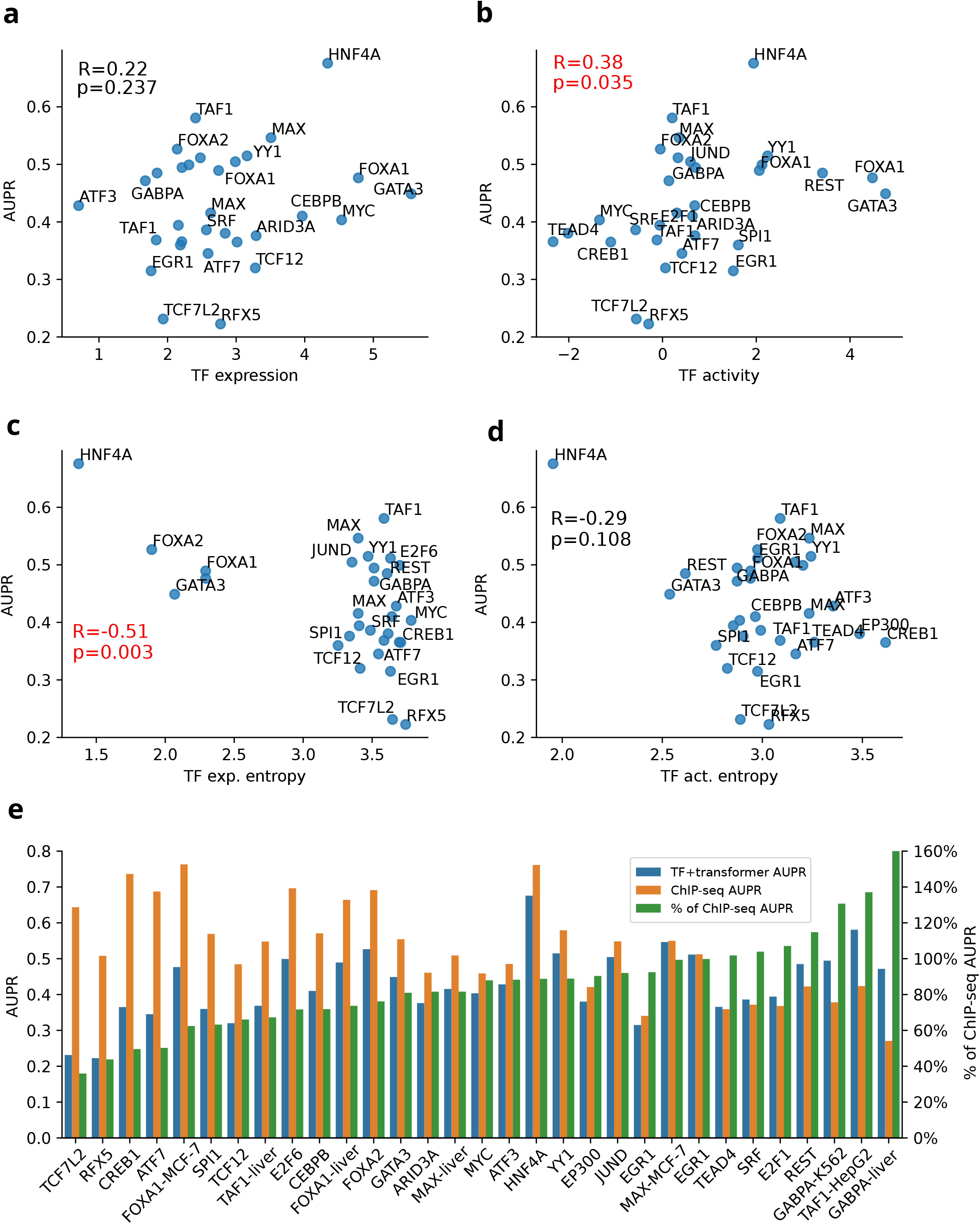
Determinants of model prediction performance. a-d) Scatterplots illustrating the relationship between model performance and TF expression, TF activity, TF expression entropy and TF activity entropy. Pearson correlation coefficients (R) and p-values are shown, with significant correlations marked in red. e) Comparison of model performance against predictions based on average ChIP-seq signal. Green bars represent the ratio of model performance to ChIP-seq performance.

We hypothesized that the predictability of TF binding might differ between tissue-specific and broadly expressed TFs. Tissue-specific TFs can afford to have simpler rules regarding DNA binding preferences, as they can simply bind to accessible TFBS in their native tissues. In contrast, constitutively expressed TFs likely require more complex rulesets, making their binding harder to predict by machine learning models. To test this hypothesis, we quantified tissue specificity using expression entropy, calculated across the expression profiles of TFs in 80 different conditions. Low entropy indicates tissue-specific expression, whereas high entropy reflects a more uniform, ubiquitous expression. Indeed, our results revealed a strong negative correlation between expression entropy and model performance (Pearson’s R = -0.51, p < 0.01), supporting that tissue-specific TFs are easier to predict than broadly expressed TFs (Fig. 5c, d). It should be noted that CTCF is an exception to this trend, as it exhibits both constitutive binding and expression, and also has a very well-defined binding motif. These characteristics allow for highly accurate prediction of CTCF binding, despite its broad expression profile.

In order to account for experimental noise and the limitations of binary ChIP-seq binding labels, we also assessed performance using mean ChIP-seq signal (fold change over control). Ideally, the AUPR of using quantitative ChIP-seq signal to predict ChIP-seq labels should approach 1. However, due to the signal-noise ratio of the experiment, the technical noise from peak-calling and the fixed genomic coordinates, which may not perfectly align with ChIP-seq peaks, the actual performance is reduced.

Our analysis showed that AUPRs using ChIP-seq signal ranged from 0.76 to as low as 0.27 for some TFs. For example, GABPA in liver achieved an AUPR of only 0.27, whereas our TF+transformer model reached an AUPR of 0.47, exceeding the ChIP-seq performance by more than 70%. Another example is EGR1, where both ChIP-seq and the TF+transformer model yielded similar AUPRs of 0.34 and 0.31, respectively. Performances in different TFs can be normalized in this manner relative to the ChIP-seq prediction performance. This metric helps to highlight TFs where there is room for improvement. For instance, TCF7L2 exhibited the lowest relative performance, with a TF+transformer model AUPR of 0.23 compared to 0.64 for ChIP-seq, representing only 36% of the ChIP-seq level. On the other hand, despite similar absolute AUPR scores for EGR1 and ATF7 (0.31 and 0.34, respectively), their relative performances diverged: EGR1 achieved 92% of the ChIP-seq performance, while ATF7 only reached 50%. This suggests that improving ATF7 predictions may be easier than improving EGR1 predictions, as the latter is already nearing the upper bound imposed by ChIP-seq performance (Fig. 5e).

### Predicting previously-unseen TFs

To systematically evaluate the performance of our models on previously-unseen TFs, we curated a new ChIP-seq dataset containing NR2F2, ZBTB33, RXRA, RAD21, and HNF4G in liver tissue and ISL1, TFAP2B, FOSL2, NFIC, and PBX3 in the SK-N-SH neuroblastoma cell line. The only TF-related information the models have access to are their binding motif PWMs and their mRNA expression. The predictions we make on these TFs are simultaneously cross-TF, cross-cell type and cross-chromosome, representing the most challenging prediction scenario.

We predicted the binding of each new TF using four different approaches: i) each of the previously trained TF+transformer models, ii) the general model, iii) a newly-trained TF+transformer model trained on the new TF data, and iv) mean ChIP-seq fold change signal (Fig. 6a). The mean performance of our TF+transformer models lies between 0.15 (predicting ISL1) and 0.48 (RAD21). Unsurprisingly, newly trained TF+transformer models outperformed pre-trained models in all cases, and using the mean ChIP-seq signal was the best predictor.

**Fig 6:**
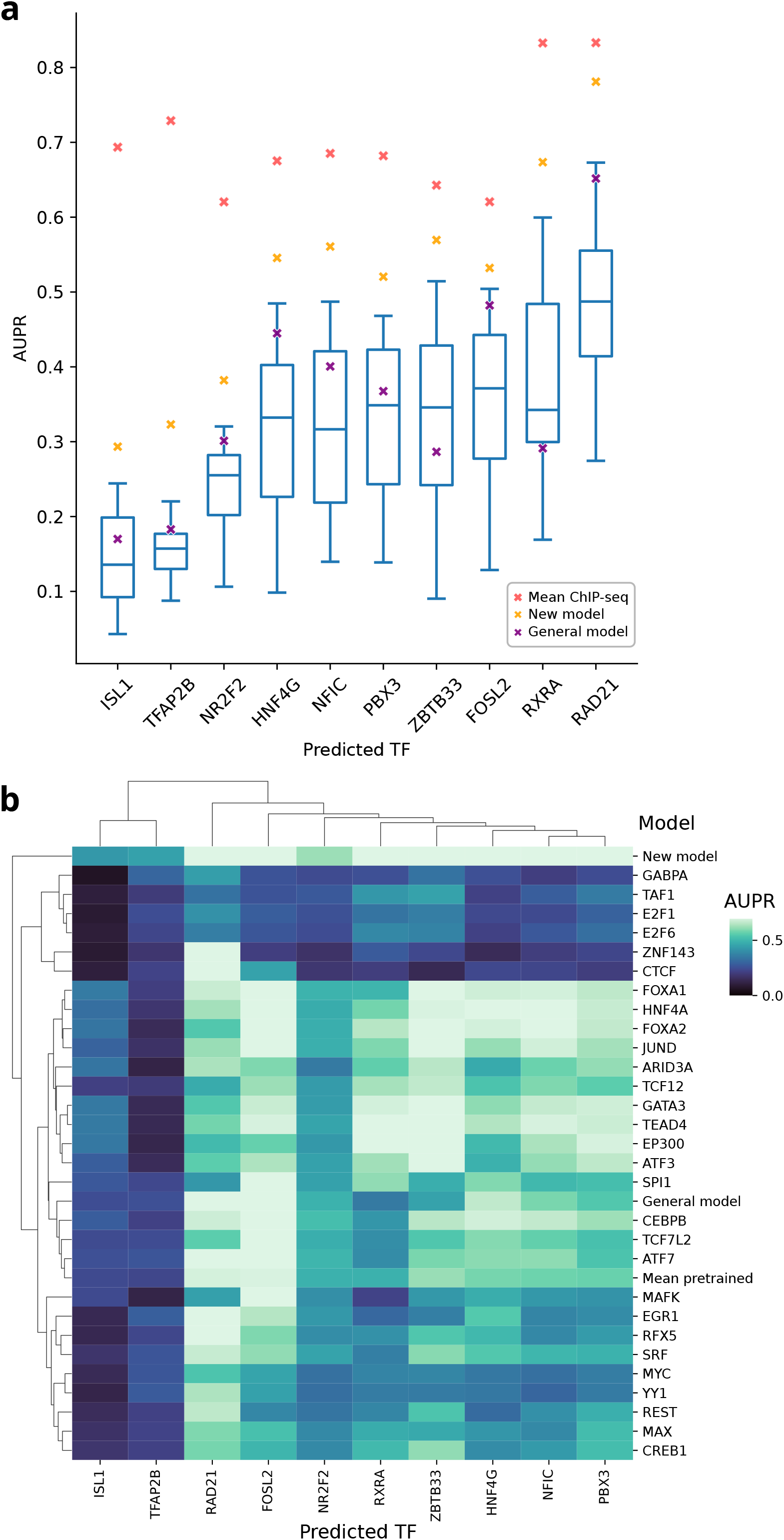
Predicting binding of previously unseen TFs. a)Prediction performance for 10 newly introduced TFs using pre-trained TF+transformer models. Yellow crosses denote the performance of models trained on data from the new TF, purple crosses denote the performance of the general model, and red crosses represent the performance based on average ChIP-seq signal (n=29 for each new TF). b) Heatmap and dendrogram showing the prediction performance of each predicted TF-model TF pair.

We observed that some of the TF models are good predictors of a new TF, while ineffective at predicting others. For example, when predicting ZBTB33 binding, the GATA3 model was the best performing TF model, having an AUPR of 0.51 —nearly matching the performance of the newly trained ZBTB33 model (AUPR 0.57). However, the GATA3 model is one of the worst predictors when predicting TFAP2B. Conversely, GABPA is the best TF model for TFAP2B, but it has poor performance in ZBTB33 (Fig. 6b). We were unable to determine a consistent rule for selecting the best TF-specific model for a newly introduced TF. Neither the similarity of expression profiles of the two TFs, nor the similarity in their binding motifs have a correlation with model performance, the latter being consistent with previous reports^30^.

Analyzing all combinations of model TFs and predicted TFs revealed distinct patterns. ISL1 and TFAP2B are challenging to predict, regardless of the model used. Furthermore, models fine-tuned on GABPA, TAF1, E2F1, E2F6, ZNF143 and CTCF consistently underperformed when predicting previously-unseen TFs. Interestingly, the CTCF model can make good predictions only on RAD21, a member of the cohesin complex which cooperates with CTCF to regulate chromosome folding^31^ (Fig. 6b).

Remarkably, the general model demonstrated robust generalization to previously-unseen TFs, achieving up to 0.65 AUPR (mean 0.36 across 10 new TFs) and outperforming the median performance of the TF+transformer models in 8 out of 10 TFs (Fig. 6a). Moreover, the general model also surpassed the ensemble mean prediction of the previously trained models. This state-of-the-art performance underscores the benefits of using transfer learning to integrate data across different TFs, generating a capability for broad generalization.

## Discussion

Our study highlights four main findings: (i) we introduce a novel two-stage transfer learning training approach that leverages the full training set while fine-tuning models for specific tasks, (ii) using DNA language model embeddings as independent features enhances the performance of simpler models, (iii) predicting *in vivo* TFBS is more accurate when TFs are highly active and when they have tissue-specific expression profiles, and (iv) TF-agnostic models can be generalized to make accurate cross-TF, cross-cell type and cross-chromosomal binding predictions.

Training prediction models that can generalize across different TFs and cell types is a long-standing goal in computational biology. On paper, such models enjoy an advantage in bigger training sets, the potential to yield deeper biological insights and of course, being able to make predictions on previously-unseen samples. However, despite the potential appeal, this approach is not commonly used in previous work. Instead, most prediction models use data from a single TF at a time. Here, we observed that using a transfer learning approach enhances prediction performance when compared to single-TF models. Recently, transfer learning was used to improve performance of a convolutional neural network to predict TF binding sites^32^. However, unlike the previous approach, our method can make predictions in cell types that were not part of the training set, which is crucial for real-world applications.

Features that are most important for TF binding prediction are DNA accessibility and sequence affinity to TF binding motifs. Other features such as enhancer probability score, TF activity, ReMap score or conservation score all contribute slightly. The single best analysis for TF binding prediction we find to be TF footprinting analysis. This is not surprising because it is in essence a hybrid feature very close to the underlying biology, since it combines accessibility information with TF binding site information, while also taking the shape of ATAC-seq coverage into account for increased accuracy.

Furthermore, our results show that the general TF binding prediction model makes accurate predictions in previously-unseen TFs, outperforming the ensemble of pre-trained TF-specific models. We extend the findings of Liu et al.^30^ on model generalization, by incorporating more powerful and accurate learning algorithms and analyzing model selection strategies. Moreover, we enhanced predictive performance by integrating sequence embeddings from a DNA language model as additional features, which significantly boosts the model’s capacity to capture complex sequence motifs.

In conclusion, computational prediction of TF binding has made significant progress in the recent decades and can provide increasingly reliable estimates of TF binding. With the growing availability of data, such as the ReMap database with binding data on 1200 TFs, transfer learning approaches present a powerful opportunity to leverage this vast source of information and develop highly accurate prediction models. Additionally, the integration of foundation models such as DNA language models enables capturing higher-level sequence features, which provide higher accuracies. With these developments, computational prediction performance is getting closer to the accuracy of ChIP-seq experiments, even though it is not yet fully capable of matching them. Models are more accurate if the TF in question is highly active in the probed condition, and when the TF is tissue-specifically expressed. Especially in these cases computational predictions can be a good alternative to experiments, if material or labor costs of the experiment are prohibitive. Looking ahead, accurate binding predictions for all TFs would be very beneficial for large scale screening applications and mapping gene regulatory networks. The development of a unified model of transcription factor binding that can make experiment-level predictions across all conditions and transcription factors would be a major breakthrough with wide-ranging implications for the field.

## Methods

### Data sources

We use the following TF-or cell-type-specific data to train our models:

1. Ground truth TF ChIP-seq labels (DREAM challenge)^33^
2. ATAC-seq data from each cell type (ENCODE)^26^
3. RNA-seq data from each cell type (ENCODE)^26^
4. Precomputed enhancer activities for 80 ENCODE cell lines and tissues^28^

a. Uses H3K27ac, H3K4me and H3K4me3 ChIP-seq data

5. TF motifs from TRANSFAC^24^ version 2022, 1 motif per TF

Additionally, we use the following data as annotation, as they do not change depending on the TF or the cell type in question:

1. ENCODE cCREs^26^, which defines our set of genomic regions
2. The hg38 human reference genome
3. Average ReMap signal track^34^
4. 30 primates phastCons^35^ scores from the UCSC Genome Browser^36,37^

### Feature overview

Genomic features include the ReMap score representing the mean signal across all available human TF ChIP-seq experiments^34^, thus providing a baseline estimate of TF binding probabilities. We also used DNA language model embeddings that provide a high-level abstract representation of the DNA sequence^25^. Moving on to motif features, we calculated the score of the best-matching TFBS and also the total affinity of the TF to the sequence, to capture the cumulative impact of weak binding sites. Furthermore, we incorporated cell type specific co-operation of TFBS, inspired by the coTRaCTE method^38^. Cell-type features include DNA accessibility and enhancer probability, and the latter is precomputed from histone modification ChIP-seq tracks. Finally, cell-and-TF features include TF expression, TF activity and the number of bound TFBS calculated by TF footprinting analysis. TF activity is calculated with a regularized linear regression model that predicts enhancer probabilities from the TF affinities in the enhancer.

### Data processing and feature extraction

We selected a set of 1.5M genomic regions from the hg38 reference sequence, consisting of ∼1M ENCODE cCREs, ∼200K ReMap peaks that do not overlap with the cCREs (to enhance bound examples), and 300K random genomic regions that do not overlap with either (to enhance the number of unbound examples).

Ground truth labels for regions: For each region in our set of genomic regions, we determined the ground truth TF ChIP-seq label from the DREAM label data, corresponding to the test set. H1-hESC cell line was excluded, because no suitable ATAC-seq data was found. NANOG was excluded as a result, since it only had training data from the H1 cell line. First, selected genomic regions’ coordinates were transferred to hg19 using LiftOver (UCSC)^36^. Then, this set of coordinates was intersected with the DREAM labels using bedtools intersect, such that at least 50% of the bin was intersecting with the region. Since DREAM bins are shorter and also overlapping with each other, this yields 4 to 10 overlapping bins for each region. The bin labels were aggregated with the following rules: if at least 40% of overlapping bins are labeled “bound” (B), then the region is labeled as (B). If at least 20% is B and 30% is “ambiguous” (A), then it is labeled as (B). If there is at least one B, but less than 20% (A), it is labeled (A). If there is at least 20% (A) but no (B), it is labeled (A). Else, it is labeled “unbound” (U).

In the leaderboard set, all regions from chromosome 8 were discarded, because all bins from this chromosome were labeled as “ambiguous” in the ENCODE-DREAM challenge files.

Motif affinity and TFBS analysis: MOODS^39^ was used to predict TFBS in the regions for each TF in the dataset, yielding the list of all TFBS with a score higher than zero. From this list, the TFBS with the highest score was selected in each region as its maximum PWM score. If a region does not have any TFBS, its maximum score is set to zero. In order to calculate the total affinity of a sequence to a TF, we used the TRAP^40^ tool. TRAP calculates p-values of the hypothesis that the total affinity of a TF to a given sequence is higher than the affinity to a set of control sequences. As control, we used a set of randomly selected 30K ENCODE-ELS enhancer regions. For the model, -log(0.0001+p) was used as the final score, instead of the raw p-value.

TF footprinting: TOBIAS^14^ version 0.16.1 was used to get TF footprinting signals and bound TF predictions. We used ATACorrect with the ATAC-seq BAM files, the hg38 human reference genome, and IDR thresholded ATAC-seq peaks from ENCODE. Afterwards, the ScoreBigwig and the BINDetect tools were used, the latter with a threshold of 0.001 bound p-value.

Bulk RNA-seq gene expression from ENCODE was downloaded as gene level TPM quantifications, and for each cell type or tissue, mean TPM was calculated across all available ENCODE data. For the model, log(1+TPM) was used as the final feature.

CRUP^29^ predictions for 80 ENCODE cell lines and tissues were used as precomputed by Sara Lopez^28^.

For ATAC-seq (fold change over control), TOBIAS TF footprinting, ReMap and phastCons scores, for which we have data in .bw format, bwtool^41^ summary was used to get min, mean and max values within our genomic regions. ATAC-seq data was quantile normalized using the qnorm (https://github.com/Maarten-vd-Sande/qnorm) python package.

### Calculating TF activity

TF activity is inspired by the tool ISMARA^42^ and it is defined as the contribution of increased sequence-TF affinity to enhancer activity. We calculated TF activity by solving for W in the formula *A* × *W* = *C*, where 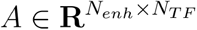 is the affinity matrix, 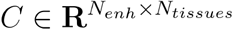 is the enhancer activity matrix, *N*_*enh*_is the number of enhancers, *N*_*TF*_ is the number of TFs, and *N*_*tissues*_ is the number of tissues. The matrix *A* is calculated by scanning enhancer sequences using MOODS with the PWMs of each TF from the TRANSFAC recommended motifs. Then, for each enhancer and TF, the score of the best TFBS in the sequence is selected as the sequence affinity to the TF motif. In cases where there are no TFBS detected in the sequence, the affinity is set to zero. The matrix *C* is the precomputed CRUP enhancer scores from Ref. 28. The TF activity matrix 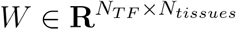 is learned through linear regression using the scikit-learn package. Afterwards,

TF activities are scaled by a factor of 10000 for easier readability, which yields the final features that are used by our models.

### Calculating cooperating TFs

This feature, inspired by the coTRaCTE^38^ framework, aims to identify TFs that bind cooperatively in a cell type specific manner. Two parameters are defined: n=5000 and k=1000. A list of ubiquitously accessible regions is created, by selecting all enhancers with a mean signal of at least s in all cell types in the training set. The value of s is set such that exactly n enhancers meet this criterion. Next, a list of cell type specific enhancers is generated for each cell type, by filtering out enhancers with very low mean accessibility, then ranking them by their relative deviation from the mean across all cell types, and selecting the top n enhancers.

The ubiquitous and cell type specific enhancers are scanned with the PWMs of all TFs, using the TRAP tool to capture total binding affinity. Then, for each TF in the training set, top k enhancers ranked by affinity to the TF are labeled as bound amongst the 2n ubiquitous and cell type specific enhancers. The same procedure is repeated separately for all TFs. Afterwards, the 2n regions are split again into ubiquitous and cell type specific groups with n enhancers each.

In the cell type specific regions, we count the number of enhancers bound by both TFs, as well as those bound by only one or neither. These counts are organized into a 2-by-2 contingency table, from which a p-value is calculated using Fisher’s exact test, to determine whether the TFs bind independently. This is repeated for the ubiquitous regions. Finally, the cooperation score L is computed as the logarithm of the ratio between the p-values for cell type specific and ubiquitous regions. If L > 2, the two TFs are considered as cooperating in this particular cell type. To increase precision, we only consider TF pairs that are bound together in at least 10 cell type specific enhancers.

For each TF in the training set, the top four cooperating TFs are selected based on their cooperation score L. For these TFs, the number of TFBS in the enhancer and the score of the best-matching site are calculated across all training enhancers, and these values are added as eight new features to the feature set.

### Model training

We implemented a multi-stage model training process using XGBoost^43^. First, the general model is trained on the complete training set, each training enhancer repeated in the set as many times as there are different TF-cell type pairs. Then, predictions from this general model were included as an additional feature in the subsequent model training stage. Further, TF-only and TF+transformer models were trained separately for each TF, using only training data corresponding to that TF.

All models were trained using XGBoost with the gradient boosting tree method. The objective function was set to logistic regression to predict TF binding probabilities. Having a probability score is necessary to rank all predictions and obtain a precision-recall score. Training parameters were set close to default values, including n_estimators=500, learning rate=0.05, subsample=0.8, column sample=0.8, tree depth=6, min_child_weight=1. For the criss-cross TF+transformer models, early stopping rounds=50 was used (see section below). The training was performed using CUDA cores.

### ^“^Criss-cross^”^ training

Our TF+transformer model uses all the previously described features, in addition to embeddings from the Nucleotide Transformer. TF+transformer models were trained only on data from a specific TF, due to memory limitations when pooling data from all TF-cell type pairs. Additionally, in order to make better predictions on previously-unseen cell types, the “criss-cross training” idea from Li et al. was used with some modifications^10^. Given a TF, we split the available training tissue types in two groups called A and B. Furthermore, we split the training chromosomes in two groups called 1 and 2. Then, we train 4 models for each TF: the first model is trained on tissue types A and chromosome group 1, while being validated on tissues B and chromosomes 2. The second model is trained on A2, validated on B1 etc. Each model training is stopped after their performance in the validation sets do not improve for 50 cycles of boosting iteration. The final score is calculated as the mean of the four models. The benefit of this approach is to reduce overfitting of the model and provide a better validation scenario, as the actual test set always is on a previously-unseen tissue or cell type.

## Code availability

The code used to extract features, train models and reproduce the figures is available at https://github.com/ekinda/tfbs_prediction_paper.

## Acknowledgements

We thank Stefan Haas for proofreading the manuscript and his helpful feedback.

## Author Contributions

E.D.A. conceptualized the study, developed the computational methods, performed data analysis and wrote the manuscript. M.V. supervised the study. All authors wrote, read, and approved the final manuscript.

## References

1. Sa, L. et al. The Human Transcription Factors. Cell 172, (2018).

2. Johnson, D. S., Mortazavi, A., Myers, R. M. & Wold, B. Genome-wide mapping of in vivo protein-DNA interactions. Science 316, 1497–1502 (2007).

3. Pique-Regi, R. et al. Accurate inference of transcription factor binding from DNA sequence and chromatin accessibility data. Genome Res. 21, 447–455 (2011).

4. Chen, X., Yu, B., Carriero, N., Silva, C. & Bonneau, R. Mocap: large-scale inference of transcription factor binding sites from chromatin accessibility. Nucleic Acids Res. 45, 4315–4329 (2017).

5. Behjati Ardakani, F., Schmidt, F. & Schulz, M. H. Predicting transcription factor binding using ensemble random forest models. F1000Research 7, 1603 (2019).

6. Fu, L. et al. Predicting transcription factor binding in single cells through deep learning. Sci. Adv. 6, eaba9031 (2020).

7. Karimzadeh, M. & Hoffman, M. M. Virtual ChIP-seq: predicting transcription factor binding by learning from the transcriptome. Genome Biol. 23, 126 (2022).

8. Qin, Q. & Feng, J. Imputation for transcription factor binding predictions based on deep learning. PLOS Comput. Biol. 13, e1005403 (2017).

9. Quang, D. & Xie, X. FactorNet: a deep learning framework for predicting cell type specific transcription factor binding from nucleotide-resolution sequential data. Methods San Diego Calif 166, 40–47 (2019).

10. Li, H., Quang, D. & Guan, Y. Anchor: trans-cell type prediction of transcription factor binding sites. Genome Res. 29, 281–292 (2019).

11. Keilwagen, J., Posch, S. & Grau, J. Accurate prediction of cell type-specific transcription factor binding. Genome Biol. 20, 9 (2019).

12. Cazares, T. A. et al. maxATAC: Genome-scale transcription-factor binding prediction from ATAC-seq with deep neural networks. PLOS Comput. Biol. 19, e1010863 (2023).

13. Li, Z. et al. Identification of transcription factor binding sites using ATAC-seq. Genome Biol. 20, 45 (2019).

14. Bentsen, M. et al. ATAC-seq footprinting unravels kinetics of transcription factor binding during zygotic genome activation. Nat. Commun. 11, 4267 (2020).

15. Aibar, S. et al. SCENIC: Single-cell regulatory network inference and clustering. Nat. Methods 14, 1083–1086 (2017).

16. Bravo González-Blas, C. et al. SCENIC+: single-cell multiomic inference of enhancers and gene regulatory networks. Nat. Methods 20, 1355–1367 (2023).

17. High-performance single-cell gene regulatory network inference at scale: the Inferelator 3.0 | Bioinformatics | Oxford Academic. https://academic.oup.com/bioinformatics/article/38/9/2519/6533443.

18. Kamal, A. et al. GRaNIE and GRaNPA: inference and evaluation of enhancer-mediated gene regulatory networks. Mol. Syst. Biol. 19, e11627 (2023).

19. Yuan, Q. & Duren, Z. Inferring gene regulatory networks from single-cell multiome data using atlas-scale external data. Nat. Biotechnol. 1–11 (2024) doi:10.1038/s41587-024-02182-7.

20. Alipanahi, B., Delong, A., Weirauch, M. T. & Frey, B. J. Predicting the sequence specificities of DNA- and RNA-binding proteins by deep learning. Nat. Biotechnol. 33, 831–838 (2015).

21. Avsec, Ž. et al. Base-resolution models of transcription-factor binding reveal soft motif syntax. Nat. Genet. 53, 354–366 (2021).

22. Avsec, Ž. et al. Effective gene expression prediction from sequence by integrating long-range interactions. Nat. Methods 18, 1196–1203 (2021).

23. Wang, K. et al. BERT-TFBS: a novel BERT-based model for predicting transcription factor binding sites by transfer learning. Brief. Bioinform. 25, bbae195 (2024).

24. Ji, Y., Zhou, Z., Liu, H. & Davuluri, R. V. DNABERT: pre-trained Bidirectional Encoder Representations from Transformers model for DNA-language in genome. Bioinformatics 37, 2112–2120 (2021).

25. Dalla-Torre, H. et al. The Nucleotide Transformer: Building and Evaluating Robust Foundation Models for Human Genomics. 2023.01.11.523679 Preprint at 10.1101/2023.01.11.523679 (2024).

26. Moore, J. E. et al. Expanded encyclopaedias of DNA elements in the human and mouse genomes. Nature 583, 699–710 (2020).

27. Matys, V. et al. TRANSFAC® and its module TRANSCompel®: transcriptional gene regulation in eukaryotes. Nucleic Acids Res. 34, D108–D110 (2006).

28. López Ruiz de Vargas, S. A comprehensive map of predicted enhancers on a large panel of human primary cells, cell lines and tissues. (2021) doi:https://repositori.upf.edu/handle/10230/49013.

29. Ramisch, A. et al. CRUP: a comprehensive framework to predict condition-specific regulatory units. Genome Biol. 20, 227 (2019).

30. Liu, S. et al. Assessing the model transferability for prediction of transcription factor binding sites based on chromatin accessibility. BMC Bioinformatics 18, 355 (2017).

31. Mach, P. et al. Cohesin and CTCF control the dynamics of chromosome folding. Nat. Genet. 54, 1907–1918 (2022).

32. Novakovsky, G., Saraswat, M., Fornes, O., Mostafavi, S. & Wasserman, W. W. Biologically relevant transfer learning improves transcription factor binding prediction. Genome Biol. 22, 280 (2021).

33. ENCODE-DREAM in vivo Transcription Factor Binding Site Prediction Challenge.

34. Hammal, F., de Langen, P., Bergon, A., Lopez, F. & Ballester, B. ReMap 2022: a database of Human, Mouse, Drosophila and Arabidopsis regulatory regions from an integrative analysis of DNA-binding sequencing experiments. Nucleic Acids Res. 50, D316–D325 (2022).

35. Siepel, A. et al. Evolutionarily conserved elements in vertebrate, insect, worm, and yeast genomes. Genome Res. 15, 1034–1050 (2005).

36. Hinrichs, A. S. et al. The UCSC Genome Browser Database: update 2006. Nucleic Acids Res. 34, D590–D598 (2006).

37. Nassar, L. R. et al. The UCSC Genome Browser database: 2023 update. Nucleic Acids Res. 51, D1188–D1195 (2023).

38. van Bömmel, A., Love, M. I.Chung, H.-R. & Vingron, M. coTRaCTE predicts co-occurring transcription factors within cell-type specific enhancers. PLoS Comput. Biol. 14, e1006372 (2018).

39. Korhonen, J., Martinmäki, P., Pizzi, C., Rastas, P. & Ukkonen, E. MOODS: fast search for position weight matrix matches in DNA sequences. Bioinformatics 25, 3181–3182 (2009).

40. Roider, H. G., Kanhere, A., Manke, T. & Vingron, M. Predicting transcription factor affinities to DNA from a biophysical model. Bioinformatics 23, 134–141 (2007).

41. Pohl, A. & Beato, M. bwtool: a tool for bigWig files. Bioinformatics 30, 1618–1619 (2014).

42. Balwierz, P. J. et al. ISMARA: automated modeling of genomic signals as a democracy of regulatory motifs. Genome Res. 24, 869–884 (2014).

43. Chen, T. & Guestrin, C. XGBoost: A Scalable Tree Boosting System. in Proceedings of the 22nd ACM SIGKDD International Conference on Knowledge Discovery and Data Mining 785–794 (Association for Computing Machinery, New York, NY, USA, 2016). doi:10.1145/2939672.2939785.

