## Supplemental Information, sup. Table 1 for "Transfer learning and DNA language models enhance transcription factor binding predictions"

27 September 2024

### Supplementary Information

**Comparability to other methods**

Due to differences between the original DREAM challenge setup and the methodology described in this work, direct comparisons between our method and those from the challenge should be approached carefully.

Firstly, there are differences in input data. We use ATAC-seq in order to be able to leverage state-of-the-art TF footprinting methods, whereas the DREAM challenge relied on DNase-seq. Additionally we use additional data types such as histone modification ChIP-seq and the average ReMap signal, which challenge contestants did not have access to. To ameliorate this advantage, we masked these features and trained "Dream-like" models, which only have a mean AUPR difference of 0.02 compared to the full model. Another distinction lies in the reference genome: we employed the hg38 build, while the challenge used hg19. To ensure consistency, we applied LiftOver to convert genomic coordinates between references.

Secondly, our analysis focused on a fixed set of 1.5M genomic regions that contain ENCODE-cCREs, randomly sampled ReMap peaks and randomly sampled negative regions. These regions range from 150 to 350 base pairs in length. In contrast, the DREAM challenge utilized the entire human genome, divided into overlapping 200 bp bins with a 50 bp sliding window. This constraint limits the training data available to our models considerably. Additionally, in the challenge, predictions were made for each bin, with multiple overlapping bins covering any given region. To compare results, we averaged prediction scores across all bins overlapping by at least 50% with our selected regions. We also tested taking the maximum score across the overlapping bins, which led to a decrease in performance and hence we excluded this from the main analysis.

The third difference lies in the design philosophy of the models. Challenge participants, such as J-Team, leveraged extensive long-range DNA accessibility features, often spanning thousands of base pairs. For example, their approach used DNase-seq data from 80 neighboring bins to predict binding in the central bin, while employing at least 37 different DNase-seq-based features. By contrast, our model is purposefully constrained to a single enhancer region (maximum 350 bp) for making TF binding predictions. This design reflects our view that the information contained within an enhancer alone should suffice to predict TF binding, as this mirrors the way binding occurs *in vivo*.

**Impact of DNA language model embeddings and criss-cross training**

As described in the Methods section of the main text, the transformer model uses all the previously described features in addition to embeddings from the genomic LLM named Nucleotide Transformer (NT). In this work by Dalla-Torre et al., the NT neural network model is trained like a regular large language model to predict masked subsequences within a larger sequence context. In order to achieve this, input one-hot encoded DNA sequences are tokenized and then data for each token is transformed into a 1024-dimensional embedding space, where they share information between them through the transformer blocks. Thus the model learns important sequence features and encapsulates abstract high-level information in the 1024-dimensional embedding. We use these embeddings as additional features in our transformer models, by taking the average embedding across all tokens of an input sequence.

The criss-cross transformer model yields slight overall improvement compared to regular transformer models: the biggest improvement is seen in MAX, improving the AUPR from 0.38 to 0.42. The mean performance improves by 0.01 AUPR, from 0.5 to 0.51, while the median stays the same at 0.49. Interestingly, we observed stark differences in model performance when comparing single models trained on the tissue types A vs B. For some TFs, training on some tissues resulted in very poor performance. While taking the mean of all four models ameliorates this, in those cases the single model can outperform the final model. We did not observe any such effect with regards to chosen chromosome groups, indicating that our models do not “memorize” chromosomal features but can be generalized to previously unseen genomic regions. Since the overall performance is better, we report performances of the transformer model with criss-cross training.

The transformer model performs very well, reaching an average AUC-PR of 0.51, a significant increase over the average of the TF-tuned models (0.48). Performance for individual TFs shows big improvements in some TFs: CTCF performance was improved from 0.7 to 0.75 in the PC3 cell line, and from 0.63 to 0.67 in iPSCs. Performance in HNF4A was improved from 0.63 to 0.68.

This improvement shows that our previous models were underfit in terms of DNA sequence. This is not surprising, because the only sequence-based features we used were all based on known TF binding motifs, either from the TF in question or from cooperating TFs from the Cotracte analysis. Using the pretrained transformer model directly, and putting the output into existing decision tree-based prediction models is a very simple and cheap way of including DNA sequence information into biological models.

Comparing the performances of all four of our models, there were 7 TFs where the TF-tuned model outperforms the TF-only model. Using the transformer model improves the performance in all of these cases, and has the best performance in five out of the seven cases. So the transformer model is similar to the “TF-only” approach as it uses only data from one TF, but nevertheless it can fix the problems the original TF-only model has.

Some special consideration should be made with regards to held-out chromosomes and the Nucleotide Transformer embeddings. Since the Nucleotide Transformer was trained on whole genome data, it includes information from the held-out chromosomes. It is not possible to assess the impact of this potential information leak easily, nor is it possible to retrain the nucleotide transformer model using only training chromosomes. This would incur considerable computational cost in training, and homologs of all sequences of human held-out chromosomes would need to be identified and held out in all the 850 different genomes used in this work, which is clearly unfeasible. However, in our testing, we found similar performance across training and test chromosomes, ensuring that this does not create an unfair advantage in practice.

### Supplementary Table 1

List of all features used in our models

| Number | Name | Depends on | | |
|---|---|---|---|---|
| | | Region | TF | Cell type |
| 1 | Maximum TFBS score | x | x | |
| 2 | Total TFBS affinity | x | x | |
| 3 | Average phastCons | x | | |
| 4 | Average ReMap | x | | |
| 5 | Average CRUP score | x | | x |
| 6 | Mean CRUP score of region across all cell types | x | | |
| 7 | Difference between (5) and (6) | x | | x |
| 8 | TF expression as average TPM in cell line | | x | x |
| 9 | TF activity | | x | x |
| 10 | TF activity - CRUP correlation p-value | x | x | x |
| 11 | TF activity - CRUP correlation coefficient | x | x | x |
| 12 | Minimum ATAC signal from region | x | | x |
| 13 | Mean ATAC signal from region | x | | x |
| 14 | Maximum ATAC signal from region | x | | x |
| 15 | Delta (12) (compared to mean across all samples) | x | | x |
| 16 | Delta (13) | x | | x |
| 17 | Delta (14) | x | | x |
| 18 | Average of (13) across all samples | x | | |
| 19 | Average TOBIAS score in region | x | | x |

| | | | | |
|---|---|---|---|---|
| 20 | Average of (19) across all samples | x | | |
| 21 | Difference between (20) and (19) | x | | x |
| 22 | Number of “bound” TFBS calls from TOBIAS | x | x | x |
| 23 | Maximum TFBS score in sequence, using Cotracte TF #1 | x | x | x |
| 24 | Maximum TFBS score, Cotracte TF #2 | x | x | x |
| 25 | Maximum TFBS score, Cotracte TF #3 | x | x | x |
| 26 | Maximum TFBS score, Cotracte TF #4 | x | x | x |
| 27 | Number of TFBS in region, using Cotracte TF #1 | x | x | x |
| 28 | Number of TFBS in region, using Cotracte TF #2 | x | x | x |
| 29 | Number of TFBS in region, using Cotracte TF #3 | x | x | x |
| 30 | Number of TFBS in region, using Cotracte TF #4 | x | x | x |
| 31 | Prediction from the general model (only in TF-tuned) | x | x | x |
| 32-1055 | Nucleotide Transformer embeddings (only in transformer) | x | | |

### Supplementary Table 2

Data table with ENCODE IDs
