## Supplemental Figures for "Transfer learning and DNA language models enhance transcription factor binding predictions"

8 November 2024

### Supplementary Figures

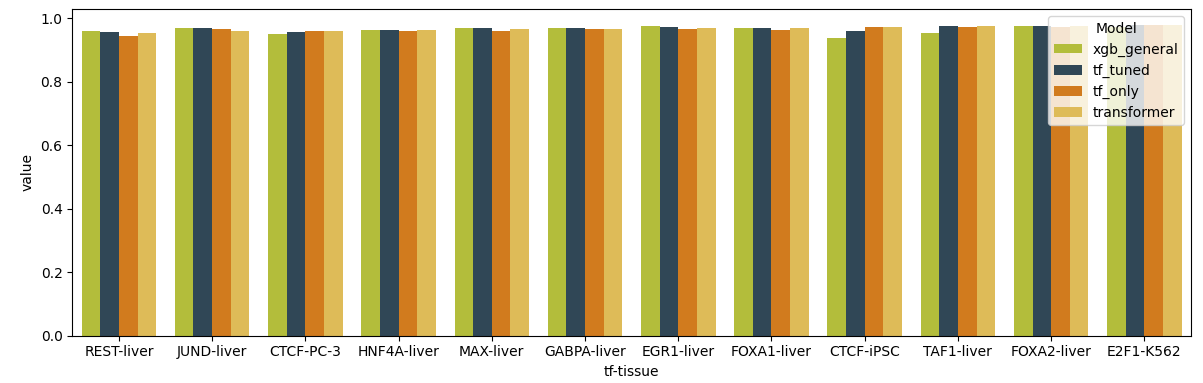

**Supplementary Fig. 1: AUROC in the test set**

Area under the reciever-operating characteristic curve (AUROC) in the test set. Due to the unbalanced nature of the dataset, AUROC is not a suitable evaluation metric.

**a**

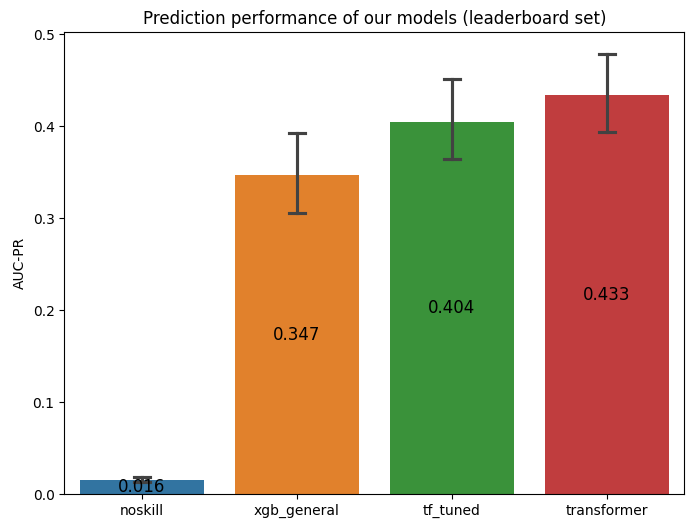

**b**

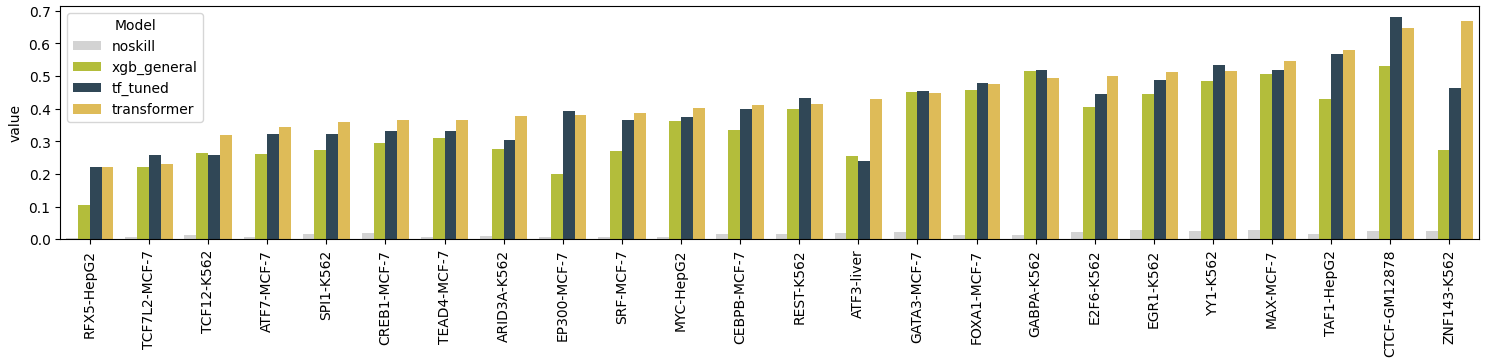

**Supplementary Fig. 2: Performance on the leaderboard set**

a) Summary of the AUPR scores in the leaderboard set of the ENCODE-DREAM challenge. The bars show the mean performance across all TFs.

b) Performance across different samples in the leaderboard set. In ATF3-liver and ZNF143-K562 there is a marked increase in accuracy using the TF+transformer model.

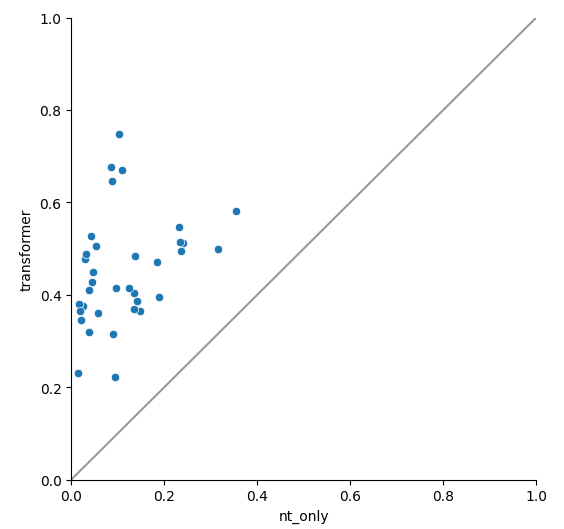

**Supplementary Fig. 3: Transformer model vs embedding-only model**

AUPR comparison in the test and leaderboard sets. Our TF+transformer model consistently outperforms models trained only using Nucleotide Transformer embeddings.
